# A Robust Chemical-free Platform for Age-Synchronized *Caenorhabditis elegans* Populations Maintenance for High-Throughput Screening in Aging Studies

**DOI:** 10.64898/2026.01.27.702075

**Authors:** Fatemeh Yousefsaber, Morteza Sarparast, Brian Johnson, Jamie Alan, Kin Sing Stephen Lee

## Abstract

*Caenorhabditis elegans* (*C. elegans*) is a well-established model for investigating the mechanisms of aging and age-related disorders like neurodegeneration. However, maintaining age-synchronized populations of *C. elegans* for aging studies without genetic or pharmacological interventions presents significant challenges, given their high reproductive rate: each nematode produces over 300 progeny. The traditional method for maintaining an age-synchronized population for multi-day studies without interventions is labor-intensive and low-throughput, hindering research on aging mechanisms and the identification of novel interventions for aging.

To address these limitations, a novel, robust method was developed to sustain age-synchronized populations in a 96-well plate liquid culture format for up to 12 days without custom-made apparatuses. The robustness of this method was substantially improved by optimizing the surface composition of the multi-well plate and disposables, refining culture parameters, including life stage, medium composition, and bacterial food concentration. To facilitate unbiased phenotype assessment throughout the lifespan, we used a Wmicrotracker ONE reader to monitor worm movement and viability in a multi-well plate. The overall fitness decline with aging using our method is comparable to that of worms maintained on solid agar. Lastly, using transgenic *C. elegans* carrying tauopathy, we demonstrated the ability of applying our optimized platform for high-throughput screening with a Z-factor of 0.7.

Our novel method simplifies age-synchronized population maintenance, enhances progeny separation, and reduces costs, enabling high-throughput screening of compounds and RNAi libraries. These advancements greatly enhance the versatility of *C. elegans* as a model organism, offering a scalable platform for genetic and compound screening and for comprehensive investigations into drug discovery and disease mechanisms.

## Introduction

Aging is a major risk factor for many chronic and degenerative diseases, including neurodegeneration, cancer, cardiovascular disease, and metabolic disorders (1). As the population aged 65 and older will triple by 2050, the social and economic burden of age-related illnesses will increase dramatically (2). Thus, uncovering the molecular mechanisms of aging—and identifying factors and interventions that modulate the pathological effects of aging—remains critical for both basic and medical research. Here, we aim to develop a high-throughput screening pipeline to identify compounds and genes that modulate aging, thereby improving our understanding of the aging mechanism and identifying novel compounds that enhance the health span of the elderly (3,4).

Model organisms play a pivotal role in illuminating conserved pathways, and among them, *Caenorhabditis elegans* (*C. elegans*) is a well-established model for discovering novel biological phenomena within the conserved biological pathways including those recognized by the Nobel Prize for various reasons (5–8). The short lifespan, transparent body, well-annotated genome, and amenability to genetic and pharmacological manipulation make it exceptionally well-suited for high-throughput, discovery-driven research. Despite these advantages, the lack of reliable, scalable platforms that enable screening on intact, age-synchronized adult worm populations limits their use in mechanistic insights into aging pathways and pharmaceutical interventions with healthy aging.

Numerous methods have been developed for performing HT screening with nematodes in liquid media (13–16). However, these approaches rely heavily on chemical agents, such as 5-fluoro-2’-deoxyuridine (FUdR), to maintain age-synchronized populations without contamination by hundreds of progeny. Notably, FUdR has been shown to adversely affect mitochondrial function and phenotypic outcomes, such as nematode size and DNA integrity, significantly impacting the outcomes of many aging studies and hindering reproducibility across laboratories (13–15). While PDMS-based microfluidic devices have been developed for HT screening without the use of FuDR, they are incompatible with lipophilic compounds which are commonly found in drug candidates in the pipeline. In addition, these devices require specialized fabrication, prohibiting other researchers from using these designs (16–19).

In this study, we developed a novel platform for maintaining age-synchronized populations of *C. elegans* in liquid culture over an extended period for HT screening, without any chemicals or genetic interventions. This protocol starts with Day 1 adult worms that have been pre-cultured on solid nematode growth medium (NGM). Unlike traditional methods, this approach does not require bleaching, avoiding potential bleaching-induced phenotyping changes, and supports both viable and non-viable food sources for worms, enabling genetic and compound screening within a carefully controlled environment (20).

Our results demonstrated that our robust platform, with a Z-factor exceeding 0.7, can be used for HT screening of chemical libraries and potentially for genomic studies focusing on adult *C. elegans* populations. Its scalability and adaptability render it a robust alternative to chemical-dependent age-synchronization techniques, with significant implications for research on aging and related disciplines.

## Method and materials

### General experimental parameters

All subsequent liquid culture experiments described in this study use day 1 adult worms unless specified in the experiment procedures. Preparing this synchronized population requires specific steps involving the selected food source and essential reagents. The strains and general supplies are listed below:

## Key resources table

| Reagent or Resource | Source | Identifier |
| --- | --- | --- |
| <u>Bacterial strains</u> |  |  |
| *E. coli* OP50 | CGC | *E. coli* OP50 |
| *E. coli* TOP10 | Thermo Fisher Scientific | *E. coli* TOP10 |
| <u>C. elegans Strains</u> |  |  |
| N2 Bristol | Caenorhabditis Genetics Center | N2 |
| CK1441 (*Paex-3::Tau WT* (4R1N) ; *Pmyo-2::dsRED*) | A gift from Nicole Liachko | N/A |
| <u>Chemicals</u> |  |  |
| Cholesterol | Alfa Aesar | CAS: 57-88-5 |
| Agar | Fisher Bioreagents | CAS: 9002-18-0 |
| Tryptone | Fisher Bioreagents | CAS: 91079-40-2 |
| Bacto Tryptone | Gibso | REF# 212750 |
| Bacto Yeast Extract | Gibso | REF# 211705 |
| Sodium Chloride | VWR chemical | CAS: 7647-14-5 |
| Magnesium Sulfate heptahydrate | Fisher Chemical | CAS: 7791-18-6 |
| Potassium Phosphate, monobasic | Sigma Aldrich | CAS: 7778-77-0 |
| Potassium Phosphate, dibasic, | Research product international | CAS: 7758-11-4 |
| Calcium Chloride (anhydrous) | Fisher Chemical | CAS: 10035-04-8 |
| Disodium EDTA | Fisher Scientific | CAS: 64-19-7 |
| FeSO<sub>4</sub>. 7H<sub>2</sub>O | Sigma Aldrich | Cas: 7782-63-0 |
| MnCl<sub>2</sub>. 4H<sub>2</sub>O | Sigma Aldrich | Cas: 13446-34-9 |
| ZnSO<sub>4</sub>. 7H<sub>2</sub>O | Sigma | Cas: 7446-20-0 |
| CuSO<sub>4</sub>. 5H<sub>2</sub>O | ACROS ORGANICS | Cas: 7758-98-7 |
| Potassium citrate | Sigma Lifescience | 6100-05-6 |
| Ethanol | Koptec | CAS: 9002-18-0 |
| Other |  |  |
| Clear flat bottom polystyrene<br>96-well plate | Merck Milipore Ltd | CAT#: MATRNPS50 |
| Clear flat bottom Ultra-low<br>attachment polystyrene 96<br>well plate with Lid | CORNING, Costar® | REF: 3474 |
| 20 µm Nylon Mesh | Merck Milipore Ltd | REF: MANMN2010 |
| Sterile Filter Durapore™<br>PVDF Membrane, 0.22 µm | Merck Milipore Ltd | REF: SLGUR33RS |
| Cell strainer (40 µm) | MTCBio incorporated | C4040 |
| 200 µL Pipette tips | Dot scientific inc. | RY0200-BP |

The following sections outline detailed protocols for this preparation process, including necessary materials and procedures for each step.

### Preparation of essential buffers and solutions

#### 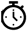 Timing: 1 day

The buffers and solutions used for this method are phosphate-buffered saline (PBS), S-basal, Trace metal solution and S-complex. All solutions must be autoclaved before use in any experiments. The details on reagents and procedures are as follows.

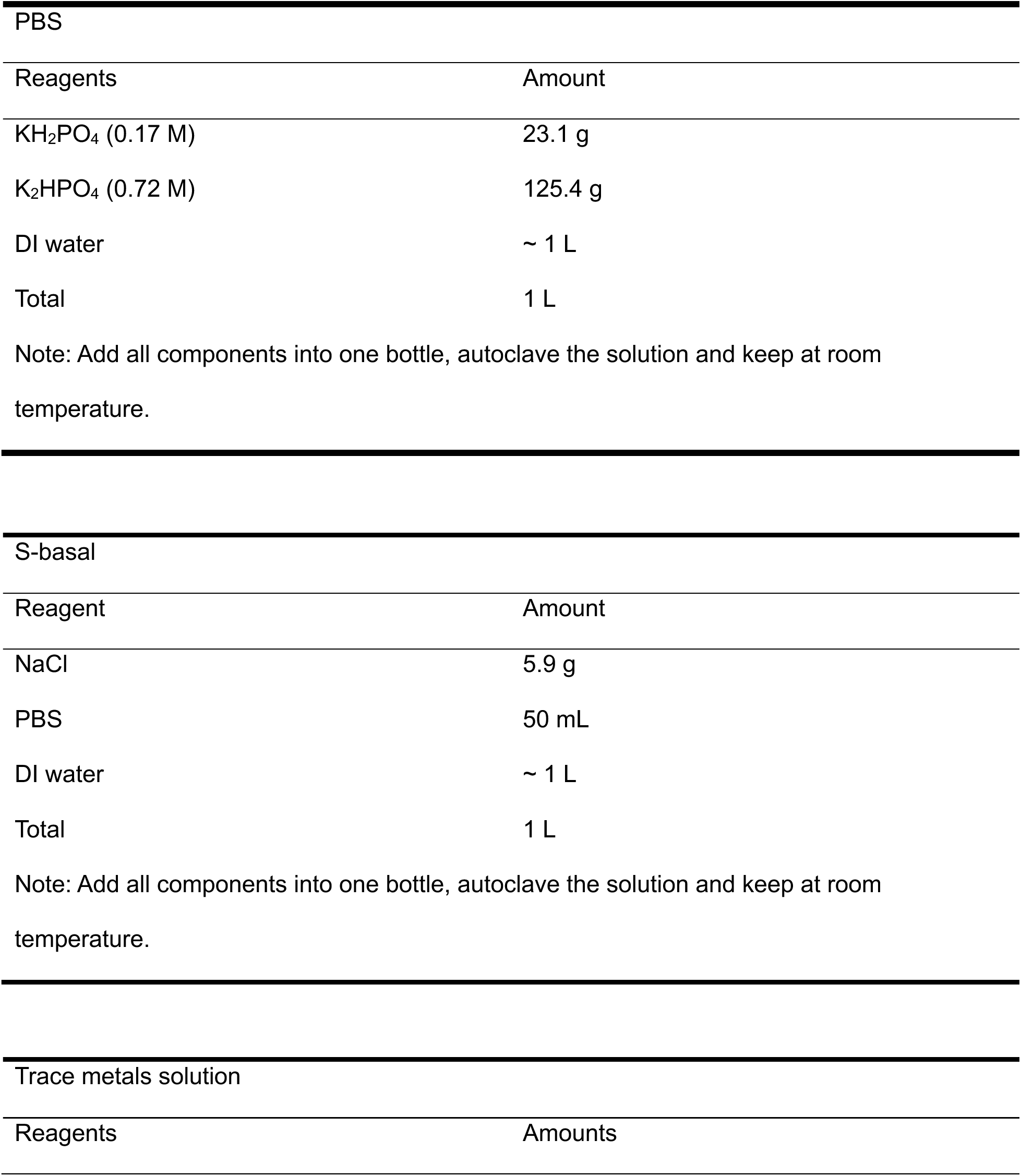
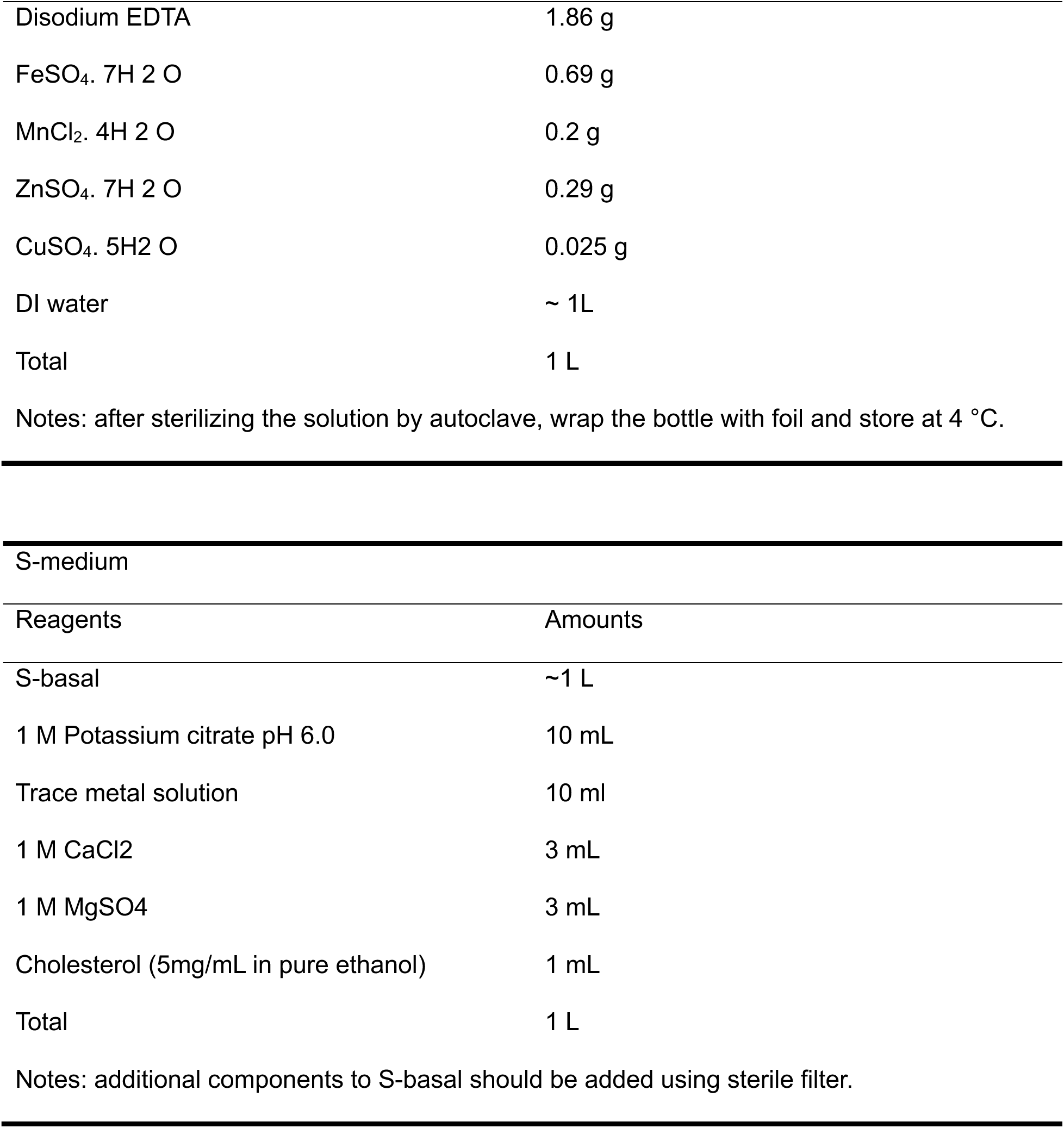

| PBS |  |
| --- | --- |
| Reagents | Amount |
| KH<sub>2</sub>PO<sub>4</sub> (0.17 M) | 23.1 g |
| K<sub>2</sub>HPO<sub>4</sub> (0.72 M) | 125.4 g |
| DI water | ~ 1 L |
| Total | 1 L |

Note: Add all components into one bottle, autoclave the solution and keep at room temperature.

| S-basal |  |
| --- | --- |
| Reagent | Amount |
| NaCl | 5.9 g |
| PBS | 50 mL |
| DI water | ~ 1 L |
| Total | 1 L |

Note: Add all components into one bottle, autoclave the solution and keep at room temperature.

| Trace metals solution |  |
| --- | --- |
| Reagents | Amounts |
| Disodium EDTA | 1.86 g |
| FeSO<sub>4</sub>. 7H<sub>2</sub>O | 0.69 g |
| MnCl<sub>2</sub>. 4H<sub>2</sub>O | 0.2 g |
| ZnSO<sub>4</sub>. 7H<sub>2</sub>O | 0.29 g |
| CuSO<sub>4</sub>. 5H<sub>2</sub>O | 0.025 g |
| DI water | ~ 1L |
| Total | 1 L |

Notes: after sterilizing the solution by autoclave, wrap the bottle with foil and store at 4 °C.

S-medium
| Reagents | Amounts |
| --- | --- |
| S-basal | ~1 L |
| 1 M Potassium citrate pH 6.0 | 10 mL |
| Trace metal solution | 10 ml |
| 1 M CaCl<sub>2</sub> | 3 mL |
| 1 M MgSO<sub>4</sub> | 3 mL |
| Cholesterol (5mg/mL in pure ethanol) | 1 mL |
| Total | 1 L |

Notes: additional components to S-basal should be added using sterile filter.

### Preparation of nematode growth medium (NGM)

#### 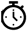 Timing: 2 days

To prepare nematode growth medium (NGM), 15 g agar, 3 g NaCl, 2.5 g tryptone, and 1 mL of 5 mg/mL cholesterol dissolved in ethanol were mixed in 25 mL phosphate-buffered saline (PBS). The mixture was further diluted into deionized (DI) water to a final volume of 1 L. The solution was autoclaved at 124 °C for 1 hour under 20 psi pressure (Consolidated Sterilizer System). After autoclaving, the solution was maintained at room temperature to cool slightly to approximately 55 °C, followed by the addition of 1 mL of 1 M MgSO_4_ and 1 mL of 1 M CaCl_2_ to achieve a final concentration of 1 mM for both MgSO_4_ and CaCl_2_. The prepared medium was poured into petri dishes (100 mm) and kept at room temperature until fully cooled and solidified. Plates remained at room temperature for 24 hours to check for contamination. NGM plates were then stored at 4 °C until use.

### Preparation of bacterial food source

#### 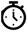 Timing: 4 days

On day 1, LB medium was prepared by dissolving 10 g of Bacto-tryptone, 5 g of Bacto-yeast extract, and 10 g of NaCl in 1 L of distilled water within a 1 L glass bottle. The solution was autoclaved for 60 minutes at 124 °C and 20 psi (Consolidated Sterilizer System), followed by storage in a dark room at room temperature for 24 hours to monitor for contamination.

### Preparation of E. coli OP50 bacterial culture

On day 2, *E. coli* OP50 bacterial culture was prepared by transferring 1 mL of the CGC-purchased stock to the prepared LB solution, followed by incubation at 37 °C for 24 hours. The culture was tested for contamination by seeding 3–5 NGM plates and inspecting them two days later. The resulting bacterial culture was stored at 4 °C prior to use.

In order to harvest the bacteria (*E. coli* OP50) fresh daily to maintain viability, sterile polypropylene test tubes (15 or 50 mL, preferably 15 mL) were weighed prior to use. The bacterial suspension was centrifuged at 7,000 rpm (Eppendorf 5430R Refrigerated Centrifuge, United States) for 10 minutes at 4 °C, the supernatant was aspirated, and fresh bacterial suspension was added; this was repeated until the entire culture volume was processed. The resulting pellet was sequentially washed with PBS up to five times, until the supernatant appeared clear. Pellets were then resuspended in S-complex medium. Prior to assay preparation, optical density at 600 nm was measured using Tecan Infinite M1000 Pro Plate Reader (Figure S1)

### Preparation of the Ghost bacterial culture

To prepare Ghost Bacteria (GB), 1 mL of *E. coli* TOP10 (already transformed with pλPR cI-Elysis, Addgene) or obtained from an existing stock added to LB medium, then incubated at 28 °C for 24 hours. The culture was tested for contamination by seeding 3–5 NGM plates and inspecting them after 2 days. Once growth was confirmed, the bottle was transferred to a 42 °C incubator for 2 hours to induce *E* gene–mediated lysis, and subsequently stored at 4 °C.

In order to harvest the bacteria (GB), sterile polypropylene test tubes (15 or 50 mL, preferably 15 mL) were weighed prior to use. The bacterial suspension was centrifuged at 7,000 rpm (Eppendorf 5430R Refrigerated Centrifuge, United States) for 10 minutes at 4 °C, the supernatant was aspirated, and fresh bacterial suspension was added; this was repeated until the entire culture volume was processed. The resulting pellet was sequentially washed with PBS up to five times, until the supernatant appeared clear. Pellets were then resuspended in S-complex medium to a final concentration of 25 mg/mL, designated as GB concentrate, and stored at 4 °C for up to 3–4 weeks. More concentrated preparations could be stored and diluted to the optimized working concentration of 25 mg/mL before experiments. Prior to assay preparation, optical density at 600 nm was measured using Tecan Infinite M1000 Pro Plate Reader (Figure S1) (21).

### Preparing non-binding pipette tips

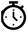 Timing: 1 day

One common challenge in carrying HT assays with *C. elegans* is the presence of hydrophobic surfaces in laboratory consumables, such as multi-well plates and pipette tips. Some researchers use glass pipettes to avoid this issue, while others use a larger population so that losing worms during the experiment won’t significantly impact the results. However, the problem of losing worms during the process, resulting in an inconsistent number of worms throughout the experiments, persists. Here, we propose a simple yet effective solution to address this issue. By applying oxygen plasma treatment to disposables like pipette tips and multi-well plates, we can enhance surface hydrophilicity and make pipette tip surfaces non-binding, facilitating precise control over *C. elegans* populations in quantitative studies. The procedure is outlined below.

To minimize losing worms through pipetting and maintaining worms in multi-well plates, we treat these consumables with oxygen plasma as outlined below:

1. The desired number of pipette tips was placed into a transparent plastic box.
2. A benchtop plasma cleaning device (Plasma Etch PE-50 Plasma etcher/cleaner, Plasma Etch, Inc, Nevada, USA) with oxygen gas was used for the surface treatment with the following settings:

1. Pressure: 250-280 mTor
2. Flow rate: 15 cc/min
3. Frequency: 100W
4. Time: 3 minutes

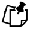 **Additional note**: Based on our experiments, plasma treatment didn’t wear off for at least 6 months. However, we do recommend testing the tips before initiating the experiment.

### Growing age-synchronized worm population

#### 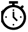 Timing: ∼3 days (strain dependent)

One key strategy to minimize experimental error is to maintain populations that are as homologous as possible, with the only variation being the factors under investigation. This is particularly crucial in developmental and aging studies, where even small differences can have significant impacts. To establish an age-synchronized population that is both healthy and well-fed, healthy adult worms were transferred onto fresh nematode growth medium (NGM) plates seeded with *E. coli* OP50 and allowed to lay eggs for approximately 24 hours. The adults were then removed, and the larvae were separated by filtration using S-basal solution and a 40 µm filter placed on top of a 50 mL centrifuge tube (22–23); adult worms were retained on top of the filter, while larvae and eggs passed through due to their smaller size. It’s critical to use new cell strainers for each new strain to minimize the risk of cross-contamination. The larvae were transferred to fresh NGM plates seeded with bacteria and maintained until they reached the last developmental stage (L4), typically within 48 hours. Once the majority of the population had reached the L4 stage, the worms were filtered again, collected from the top of the 40 µm filter, and placed on fresh NGM plates seeded with bacteria. This process was repeated to generate a new population for experiments. Once this population reaches L4 stage, they can be used for screening in liquid culture. Notably, no bleach or other chemicals were used in this method to minimize environmental factors that could impact the nematodes.

### Maintaining an age-synchronized worm population in liquid culture

#### 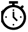 Timing: up to 12 days (strain and study dependent)

All filtrations were performed using S-basal solution, which helps balance the osmotic pressure of worms during transfer, in contrast to using DI water. Moving forward, S-basal will continue to be used as the filtering solution, while S-medium will serve as the primary culture medium for maintaining worms in liquid culture.

On day 1, the collected adult worms were dispensed in S-medium at a density of approximately 2000 worms per 1.5 mL. Using non-binding pipette tips prepared beforehand, no more than 70 worms in 50 µL with S-medium were transferred to each well of the assay plate. Images were taken with a microscope (Leica S-Series 10x/23B Widefield Adjustable Eyepiece Part # 10447137, United States attached to Flexacam C3 12 MP Microscope Camera, United States), for future data analysis, including the number of worms per well. This step was then followed by adding 10 µL of treatment solution or vehicle and an additional 20 µL of S-medium to measure the mobility score using the Wmicrotracker ONE (15 minutes, mode 01 standard average) as a post-dosing baseline. Lastly, approximately 20 µL of the desired bacterial food source at (25 mg/ml) was added to each well, resulting in a total volume of 100 µL culture medium. The lid was secured, and the plate was placed on a rotator set to 120 r/min.

On day 2, 50 µL of liquid was carefully removed from each well, keeping the pipette tip positioned on the upper side of the well to avoid disturbing the worms. The remaining liquid, containing worms, was gently transferred onto a 20 µm mesh cell strainer 96-well plate (Nylon Mesh, MilliporeSigma, Massachusetts, US). Since worms maintained in liquid culture tend to become thinner and longer, a smaller pore-size filter was used compared to the cell strainers typically used for solid culture (Figure 2C). To minimize transferring the bacterial aggregates, the pipette tip was held at the bottom of each well, opposite to the side of the bacterial aggregates. The worms on the mesh filter were further washed by adding 100 µL of S-basal to the top of the filter, placing the mesh on a dry paper towel, and gently pressing the edge of the cell strainer plate to facilitate liquid flow. This step was repeated up to five times until all bacterial aggregates were removed. While worms can be placed on ice for 10–15 minutes to filter adult worms and remove bacterial aggregates by gravity without affecting their viability (9), our observations indicated that this method was less effective than anticipated. Because the mesh filter pores are small, liquid and larvae pass through slowly; using paper towels helps speed up filtration and reduces the stress caused by prolonged exposure to filtering solution without bacterial food.

**Figure 1.**
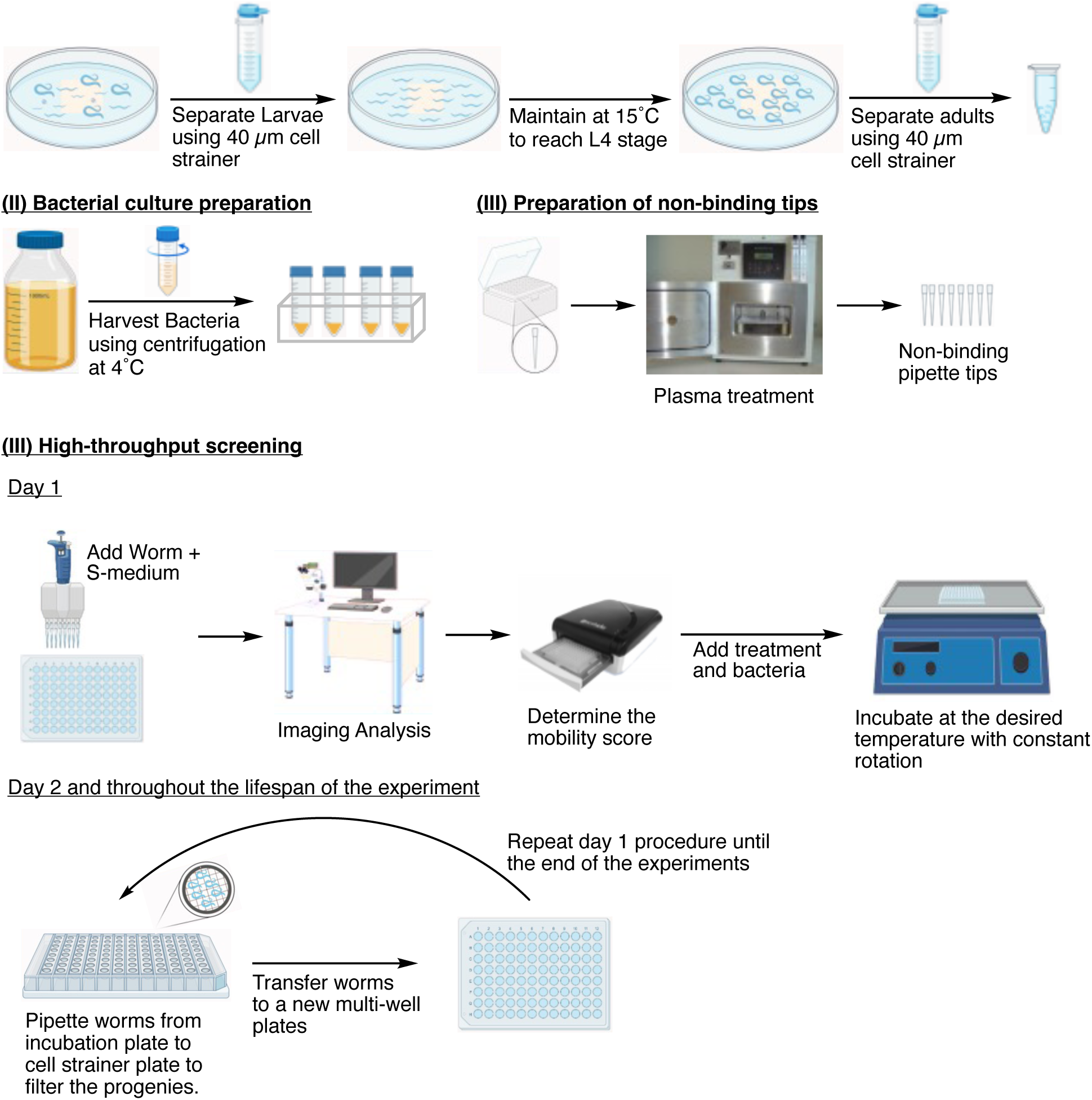
Graphical Abstract. Schematic presentation of a robust high-throughput screening platform for C. elegans.

**Figure 2.**
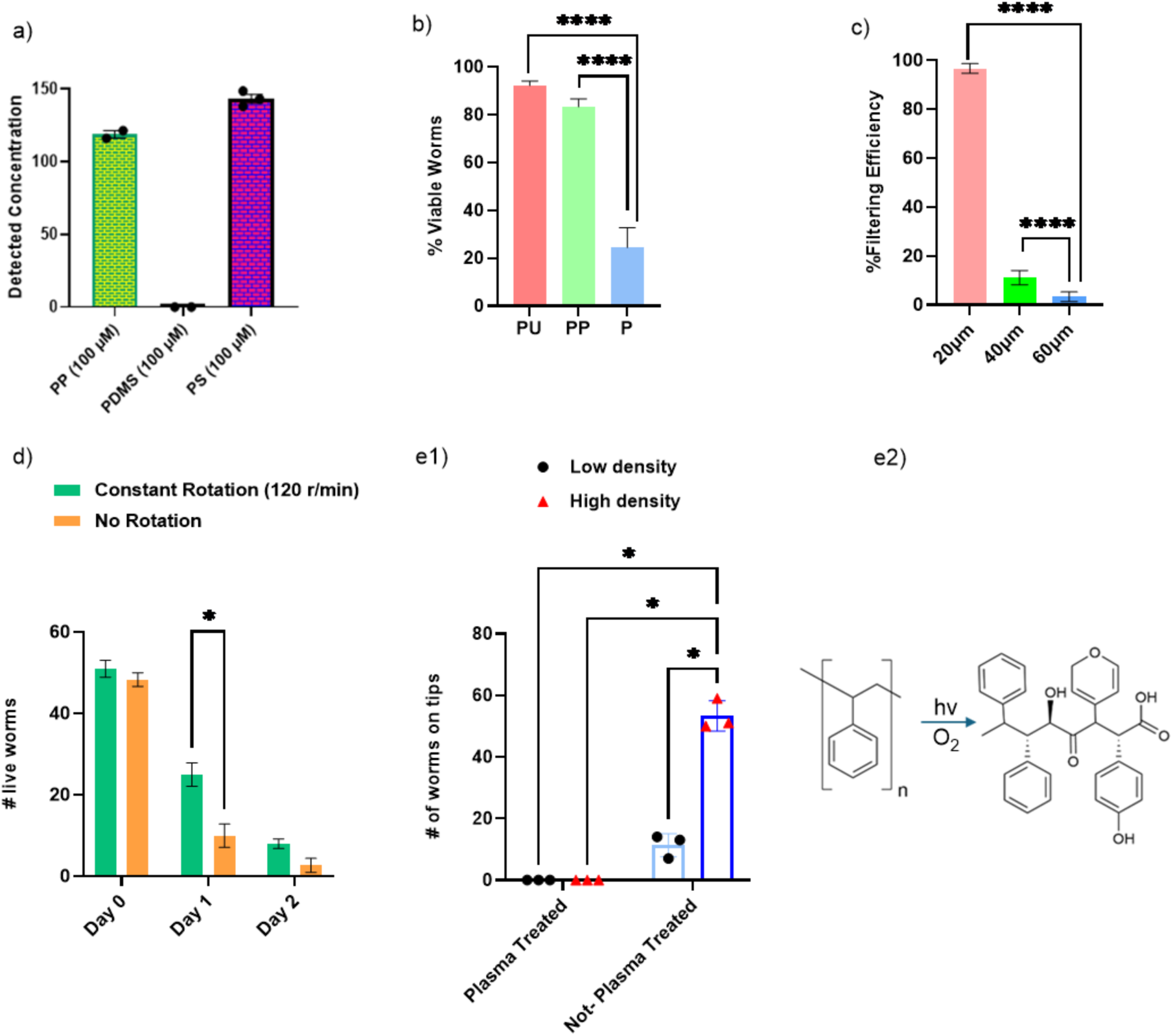
Labware surface materials impact the viability of *C. elegans* and the bioavailability of lipophilic compounds in the assays. (a) The PDMS surface significantly absorbed lipophilic lipid mediator, epoxyeicosatrienoic acid (EET), while PP and PS surfaces do not. (b) The multi-well polystyrene plate (P) decreases worm viability by ∼75%, and PU and PP plates do not have any impact on worm viability. (c) Mesh filter with a pore size of 20 µm effectively maintains adult worms, with all progenies removed. (d) Constant rotation of worms improved worm viability in liquid culture. (e1) Oxygen plasma treatment on plastic pipette tips, significantly reduces *C. elegans* attachment on the tip surface; “Low density” group refers to 500 worms in 2000 microliter solution and “High density” group refers to 500 worms in 500 microliter solution, (e2) Schematic diagram of how oxygen plasma treatment alters polystyrene surface hydrophobicity by introducing carbon–oxygen functional groups. For all experiments, N = 3, and about 20 worms were tested on each trial. For statistical analysis, a one-way analysis of variance (ANOVA) with Tukey’s post-hoc test was used. *P ≤ 0.05, **P ≤ 0.01, ***P ≤ 0.001, ****P < 0.0001.

After washing, worms retained on the filter well were resuspended in 50 µL of S-medium, and were transferred into the new assay plate. Images were taken for data analysis, baseline activity was measured using the Wmicrotracker ONE (15 minutes, mode 01 standard average), treatments were added as described before, and additional images were taken as needed. An additional 30 µL of S-medium was added, followed by a second mobility score measurement using the same settings, and approximately 20 µL of the bacterial food source was then added. Finally, the lid was placed on the plate, which was returned to the rotator (120 r/min).

Each hermaphrodite adult *C. elegans* can produce approximately 300 progeny during the first five days of adulthood (9). To maintain an age-synchronized population, adults must be separated from their offspring using the mesh filter plate. This filtration step also aids in removing bacterial clogs and ensures consistent dosing of treatments on a daily basis (Figure S3 and S4). It is critical to use new mesh filter plate when working with more than one strain to minimize the risk of cross-contamination.

### Wmicrotracker ONE setup

The Wmicrotracker ONE software was installed based on the manufacturer’s manual (see http://www.phylumtech.com for installation instructions). A new project was created based on the following parameters. The plate type was chosen based on the type of multi-well plate we used; in this article, “F96” was selected in the start window. The project and groups were labeled accordingly, and the acquisition lapse was set to 15 minutes (or longer if required). *C. elegans* was chosen as the organism, and the mode was configured to “DSP_Method_1Threshold Avg.” This method quantifies activity by analyzing the number of times organisms interrupt the infrared beam, with each interruption recorded as a unit of activity. The advanced tools option for real-time data was activated to enable re-analysis of data if needed. Once these parameters were set, the experiment was initiated, and the acquisition data were saved in the selected format.

### Image analysis

One of the key challenges in High-Throughput Screening (HTS) is maximizing efficiency while acquiring large datasets for analysis. The method outlined above, when combined with Wmicrotracker ONE technology, undoubtedly accelerates HTS processes. However, because the movement score is based on beam interference, it is crucial to determine the number of live worms in each well for data normalization (24). This becomes especially important when using different worm strains or conducting chemical, pharmaceutical, or genetic screenings, as these factors could impact worm viability and/or movement. Manual counting of live and dead worms is time-consuming and prone to bias. To address this, we recommend using image analysis software like Cell Profiler software or ImageJ. ImageJ has built-in coding modules as well as a user-friendly interface to optimize different parameters based on the type and characteristics of images. All image analysis data and instructions are included in the supporting information (Figure S8).

## Results and discussion

Despite a decade of research, the molecular mechanism of aging remains understudied, and no effective interventions have been developed. Studying the aging mechanism in whole animal models could capture the complexity of the aging process. This project aims to develop a robust HT platform that addresses limitations of current platforms to investigate the molecular mechanisms underlying age-related phenotypic changes in *C. elegans. C. elegans* has been instrumental in discovering ground-breaking biology and aging mechanisms. Specifically, our pipeline solves the long-standing challenge of maintaining an age-synchronized *C. elegans* population in a 96-well plate format without the use of chemicals or genetic manipulation, both of which could interference aging phenotypes, thereby improving the throughput of aging studies (9–12). Our pipeline was developed using commercially available equipment and supplies, without any custom-made components, such as microfluidic devices, to increase accessibility for other researchers. During development, we optimized parameters that are critical for the success of HTS with *C. elegans,* including the preparation of age-synchronized *C. elegans*, culturing conditions, preparing the consumables to minimize loss of worms during the experiments, maintenance conditions for the age-synchronized adult worm population, and phenotype measurement and analysis that are rarely discussed in detail. The optimized protocols are further discussed and described in detail below and in supplemental information, enabling other aging researchers to utilize this protocol for studying the mechanisms of aging and age-related phenotypic changes.

### Limitations of current HT assay with C. elegans

The ability to culture *C. elegans* in a liquid medium can facilitate research across various fields, including toxicology, nutritional science, developmental biology, and aging, by enabling HTS using an *in vivo* system. Culturing *C. elegans* in liquid medium dates to 1957, when Warwick Nicholas proposed the first axenic liquid cultures, which were further optimized to Habituation and Reproduction media (CeHR) by Dr. Clegg to improve growth(25–26). However, this method presented several limitations, including slow development, altered physiology, morphological changes, and susceptibility to microbial contamination. Since then, various efforts have been made to develop methodologies for using *C. elegans* in HTS to investigate biological mechanisms and bioactive compound identification, using both genetic and pharmacological means. Current methods for growing and maintaining age-synchronized populations involve bleaching a mixed population of *C. elegans* to remove adults while keeping the viable eggs. This is typically followed by using FUdR (5-fluoro-2’-deoxyuridine) to avoid progeny contamination. However, FUdR impacts phenotypes, including reduced movement, decreased pharyngeal pumping rates, abnormalities in movement patterns, body size and shape, and internal morphology, such as intestinal atrophy and degenerated gonads (27–28). Treatment with FUdR also elevates superoxide dismutase levels and extends longevity on specific transgenic worms. (29–33). González *et al.* reported that while bleaching does not affect the worm’s lifespan, at low concentrations of bleach, it results in fewer viable eggs, with changes in morphology and tissue damage observed compared to the non-bleaching method (20,34). In addition, there are other challenges to conducting *C. elegans* HTS studies, including bacterial metabolism interference in chemical screening and the difficulty of maintaining viable bacterial populations for genetic studies (35). Addressing these limitations would collectively expand the scope and precision of *C. elegans* research, contributing to its continued importance as a model organism across diverse fields of study. In this study, we have systematically developed a robust pipeline that addresses several issues in performing diverse types of HTS experiments with *C. elegans*.

### Determining the optimized surface modification for pipette tips

One of the challenges in *C. elegans* studies is transferring worms between multi-well plates for HTS. In liquid culture, worms can be transferred using either glass pipettes or plastic pipette tips. However, transferring *C. elegans* using glass pipettes is relatively low throughput. Alternatively, a multi-channel pipette is commonly used to transfer worms across multi-well plates. Yet, we found that worms tend to stick to the surface of the plastic pipette tips. This could result in a large variation in the number of worms between wells, particularly when a small number of worms is used. Various techniques can modify surface characteristics, particularly hydrophobicity, including plasma treatment and coating. Among these methods, we opted for oxygen plasma treatment due to its well-established effectiveness and ease of use compared to other coating methods, which require identifying suitable inert materials for *in vivo* studies (36). Oxygen plasma introduces various functional groups to the surface of polymers, such as carbon-oxygen groups (C–O, C═O, O═C–O, O–CO–O, and –OH) on polystyrene surfaces. These modifications significantly reduce surface hydrophobicity and enhance wettability, thereby minimizing worm adhesion. As shown in (Figure 2e and S2), worms remained fully suspended in the medium when resuspending and mixing worms with plasma-treated tips, while worms were stuck on the untreated pipette tip surface. Quantitative analysis indicated a substantial decrease in worm retention, with no worms adhering to plasma-treated tips compared to over 50% of worms adhering to untreated tips. This advancement not only resolves a common bottleneck in liquid-based *C. elegans* assays but also enhances reproducibility by ensuring consistent sample transfer and maintaining a consistent population size.

### Optimizing the surface properties of HTS labware

Polydimethylsiloxane (PDMS), polypropylene (PP), and polystyrene (PS) are polymers commonly used in biomedical devices. However, both PDMS and PP are highly hydrophobic, with contact angles of 108° and 94°, respectively (noting that smaller contact angles indicate higher hydrophilicity). These materials are prone to absorbing lipid-based compounds, whereas PS does not exhibit the same degree of absorption. Given that approximately 40% of marketed drugs and 60% of compounds in research and development are highly lipophilic, selecting appropriate laboratory utensils is crucial (37). To address this problem, we first compared lipid absorption in PDMS, PP, and PS-based multi-well plates by quantitatively measuring lipid uptake. For this, 100 µM of epoxyeicosatrienoic acid (EET), a lipid mediator with anti-inflammatory and analgesic properties, was incubated in all three types of devices (3). Analysis of the incubated solution revealed that the lipophilic EET was completely absorbed by the PDMS-based device after 24 hours of incubation at room temperature, while it remained largely intact in the PP and PS-based plates (Figure 2a).

Since PS-based plates have been widely used for HT screening, we tested their compatibility for maintaining *C. elegan*s cultures for long-term studies (38). Approximately fifty day 1 adult worms were incubated in 100 μL of S-medium with 30 μL of GB, in a PS 96-well plate at 18 °C, a standard temperature for worm maintenance. After 2 hours, the plates were examined under the microscope. Surprisingly, all the worms were dead and adhered to the bottom of the well when incubating the worms in a 96-well plate (Figure S2). To further investigate the effect of surface materials of the multi-well plate on nematode survival, we repeated the experiment using Polystyrene (P), ultra-low attachment PS (PU), and plasma-treated (PP) devices (with plasma treatment settings described in pipette tip treatment in the experimental conditions). Plasma treatment enhances surface hydrophilicity by introducing polar functional groups, creating a low-binding surface. Preparing pipette tips with plasma treatment minimizes the loss of worms by preventing worms from adhering to hydrophobic surfaces. As shown in Figure 2b, after 4 hours of incubation, the number of viable worms in the ultra-low attachment and plasma-treated devices was significantly higher compared to the untreated polystyrene plates. Approximately 90% and 80% of worms remained viable in the treated devices, respectively, whereas only 20% survived in the untreated plates. Therefore, all subsequent experiments were conducted in ultra-low attachment PS plates, as they are commercially available and easily accessible.

### Determining the optimized filter size for maintaining age-synchronized adult C. elegans

Maintaining an age-synchronized population without progeny contamination is of great importance for aging studies in *C. elegans*. The common method for growing and maintaining an age-synchronized population involves first using bleach to remove all adult worms without damaging the eggs, followed by treatment with FUdR to sterilize the resulting adult worms. However, the use of FUdR can interfere with phenotypic outcomes (14, 28). Traditionally, researchers maintain age-synchronized worms in solid agar culture by either using chemical solutions or using a cell strainer with a 40 µm pore size daily to prevent progeny contamination (10–12,14, 28, 23). These methods are labor-intensive, low-throughput, and not applicable to the 96-well plate format for HTS. To address these challenges in HTS, we propose utilizing mesh cell strainers, which are typically employed in multicellular whole-organism screenings. These filters come in various pore sizes, including 20 µm, 40 µm, and 60 µm. In liquid culture, worms tend to become thinner and longer due to increased movement compared to solid agar medium. As shown in Figure 2c and Figure S2, mesh filters with a 20 µm pore size are ideal for separating adult worms from progeny without affecting the worms’ physiology or morphology.

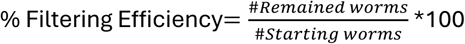

### Optimizing the culturing conditions for maintaining age-synchronized C. elegans populations

As mentioned earlier, the choice of plate material significantly impacts the maintenance of *C. elegans*. However, even with the use of ultra-low attachment PS plates, the worms survived for less than a day. According to published studies, some researchers maintain their worms with constant rotation, while others do not specify any details (11, 12). Based on our results (Figure 2d), we found that constant rotation is essential for maintaining worms in liquid culture. However, it only extended the survival period to 48 hours.

This prompts us to systematically identify factors that impact the health and viability of worms in liquid culture, which include worm life stage, culture medium components, the ratio of food to culture medium, and the volume and the food source (final food source and total volume are 20 and 100 µL respectively) . These factors were not previously discussed in detail in the literature. To ensure results without bias, worm movement was measured using the Wmicrotracker ONE, and all data were normalized to the number of worms in each well (using Image J).

#### Culture medium volume and components

One approach for conducting HTS experiments is the use of microfluidic devices, which typically hold 50 µL of total culturing medium for each well (4). Inspired by this, we first examined whether decreasing the total volume of culturing medium would extend the worms’ lifespan. To test this, approximately 50 age-synchronized day 1 adult worms were used with 30 µL GB as the food source mixed with S-medium at a total volume of 100 µL, compared to 15 µL GB mixed with S-medium at 50 µL. As shown in Figure 3a, adult worms in the 50 µL condition survived for only 3 days, whereas those in ∼100 µL survived for 6 days.

**Figure 3.**
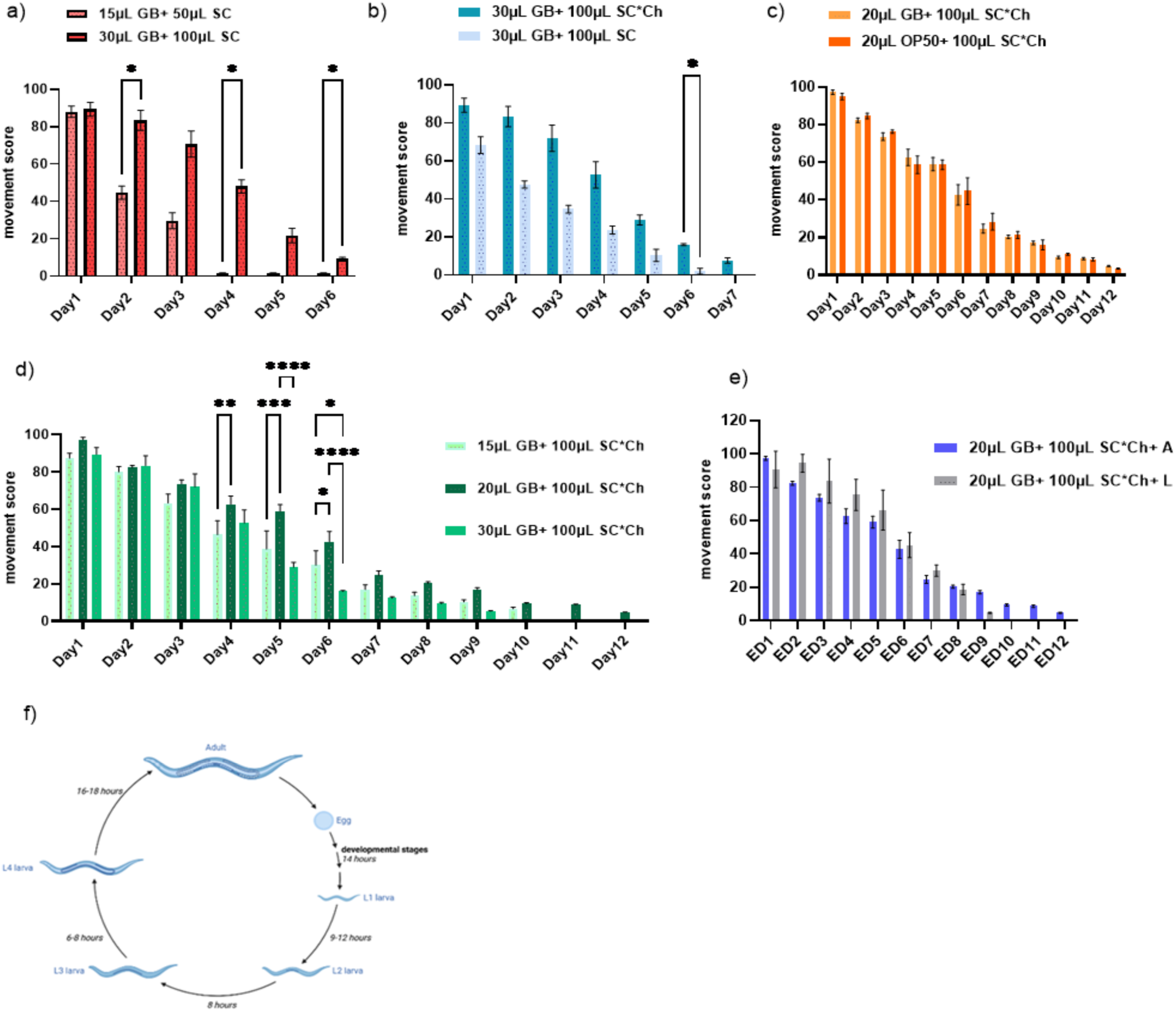
Worm maintenance conditions impact the viability of worms in liquid culture. (a) Evaluation of the effect of total liquid culture volume on worms’ survival, (b) Assessment of the effect of cholesterol on worms’ lifespan in liquid culture, (c) Comparison between viable (*E. coli* OP50) and non-viable (ghost bacteria) food sources, (d) Comparison of different food source quantities, (d) Impact of worms’ life stage on survival in liquid culture, (e) Movement comparison between adult and larvae worms in liquid culture, (f) schematic representation of *C. elegans* life cycle. . For all experiments, N = 3, and about 50 worms were tested on each trial. For statistical analysis, a two-way analysis of variance (ANOVA) with Tukey’s post-hoc test was used. *P ≤ 0.05, **P ≤ 0.01, ***P ≤ 0.001, ****P < 0.0001.

In conventional NGM culture medium, cholesterol is a key component. Since *C. elegans* cannot synthesize cholesterol *de novo*, it must acquire it from its diet. Cholesterol plays crucial roles in *C. elegans* biology, including development, reproduction, lifespan regulation, pathogen resistance, Dauer formation, intercellular signaling, and metabolic regulation. More specifically, neurons rely on *de novo* cholesterol biosynthesis for survival and growth during early development, while they depend on glial-supplied cholesterol in later stages. As a result, *C. elegans* requires dietary cholesterol to support its development (39). However, in previously published studies, cholesterol was primarily added to S-basal as a washing solution but not in a culture medium (9–12, 24). In particular, Hibshman *et al*., noted that limiting cholesterol results in calorie restriction and starvation (12). Therefore, we tested whether the addition of cholesterol impacts the health span of *C. elegans* cultured in liquid medium. As shown in (Figure 3b and Figure S5), while cholesterol extended lifespan by one additional day, no uncoordinated movement was observed, and the worms’ overall mobility score resembled those grown on an NGM agar plate. A small number of worms were lost each day during transfers to new wells.

To determine the optimal bacterial amount, a major diet for worms, specifically for *C. elegans* liquid culture, with food volumes of 15 µL, 20 µL, and 30 µL (delivered at a 25 mg/mL concentration) was tested (21). As shown in Figure 3d, the results indicate that 20 µL of food source, corresponding to a total amount of 500 µg wet weight of bacteria, provides the optimal balance. It extends the lifespan of worms while minimizing bacterial aggregates and ensuring adequate nutrient availability. Using 15 µL led to insufficient nutrition, while 30 µL resulted in excessive bacterial growth, creating unfavorable culture conditions.

### Determining the impact of food sources on C. elegans growth in liquid culture

Various food sources have been used for culturing *C. elegans* in liquid media, including CeHR medium, and various bacterial strains such as the commonly used strain *E. coli* OP50, and Ghost Bacteria (12,21,26). The choice of food source should be tailored to specific experiments. For instance, in HTS aimed at screening chemical compounds, it is best to use GB, as these non-viable bacteria prevent outcomes impacted by bacterial metabolism of the compounds. Conversely, for genetic screenings, such as RNA interference (RNAi) studies, live bacteria will be used to transfer the gene of interest. While many previous studies on *C. elegans* have been conducted using the CeHR medium, it is important to note that this medium significantly affects the experimental outcomes. Celen *et al.* reported that the effects of CeHR medium on *C. elegans* extend beyond physical changes (35), including gene regulation, reproduction, and development. In this study, we focused on GB and *E. coli* OP50 as food sources in liquid culture. As shown in Figure 3c, aligned with previous publications, no significant differences were observed between the two groups, suggesting that both food sources can be used effectively in liquid culture. However, as mentioned previously, we have further optimized the use of GB in our HTS by 1) decreasing the food source to minimize bacterial clogs, and 2) altering the transfer process to a two-step method, including a final wash in each well to separate as many worms as possible from bacterial clogs. These adjustments significantly minimize the loss of worms during the filtration process and allow us to maintain a sufficient number of worms (at least 10) in each well up to day 12 for mobility score determination. If we need to maintain the worms for the health span experiment longer than day 12, we can pool worms from multiple wells from the same treatment group to collect enough worms for mobility score testing.

### Determining the optimum conditions to maintain the C. elegans culture for aging studies

Maintaining worms in liquid culture could introduce environmental stress, which impacts their physiology. Therefore, we investigate whether starting culturing *C. elegans* at different life stages, larvae and day 1 adults, in liquid medium, impacts their health span. Larvae are smaller than adults and can pass through a 20 µm mesh filter. During the first 5 days, the culture medium was removed via a two-step aspiration to minimize the risk of losing worms (Figure S3). As shown in Figure 3e, starting liquid culture at larvae (approximately L2, L3) significantly decreased worm viability in liquid culture. Consistent with previous literature, the first batch of progeny from the larvae group appeared after 4 days of incubation in liquid culture and these progenies survived for 3 additional days (10–14). In contrast, the adult group survived until day 12. Additionally, as indicated in Figure 3e, the Larvae group at adult day 1 showed movement patterns comparable to those of day 4 and day 5 adults. These results further confirm the importance of ensuring proper development of *C. elegans* larvae in survival and behavior before any experimentation in liquid culture.

### Applying our optimized HT culturing pipeline for screening

To ensure that our pipeline does not cause any significant physiological changes to the worms, we compared the survival and mobility of *C. elegans* in liquid culture with solid agar culture. As shown in (Figure S6), no significant differences were observed between the two groups during the first 7 days. However, as the worms aged, those in liquid culture exhibited slower movement compared to their counterparts in solid agar culture. This could potentially be attributed to the constant movement of worms in liquid culture and lower food availability. Nevertheless, there was no notable difference in their physical features, such as length and width. The experiment was conducted for a maximum of 12 days, as there were not enough worms (≤10 worms per well) to measure the mobility after day 12.

The ultimate goal for this pipeline is to investigate the molecular mechanism of age-associated phenotype changes, such as neurodegeneration and identify potential interventions for these age-related conditions. Therefore, we tested our HT pipeline with transgenic worms, the Tau-Tg strain (*Paex-3::Tau WT (4R1N) ; Pmyo-2::dsRED*), as the established transgenic model for Alzheimer’s disease (38). As shown in Figures 4a and b, the Tau-Tg strain maintained in our optimized pipeline displays a significant decrease in mobility, similar to that observed in an NGM agar plate, suggesting that our platform can also be applied to transgenic strains to determine their phenotypes in HT manner.

**Figure 4.**
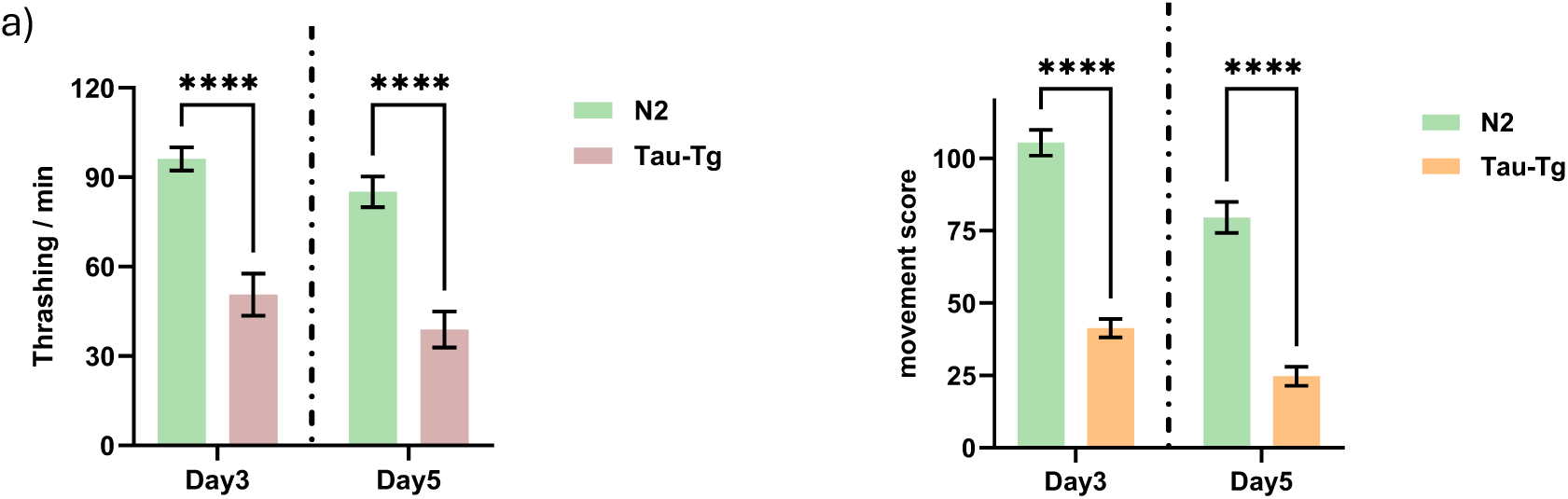
(a)Expression of Tau in neurons causes behavioral abnormalities in *C. elegans* locomotion, (b) Movement comparison between wildtype and transgenic strain in liquid culture is consistent with solid-medium culture. For all experiments, N = 3, and about ∼30 worms were tested on each trial. For statistical analysis, a two-way analysis of variance (ANOVA) with Tukey’s post-hoc test was used. *P ≤ 0.05, **P ≤ 0.01, ***P ≤ 0.001, ****P < 0.0001.

To further explore the capability of using our platform for screening against pharmaceutical interventions and RNAi to identify potential targets responsible for the observed phenotypes, we determined the z-factor of our screening platform using Tau-Tg strains and obtained a z-factor of 0.65 and 0.68 for days three and day five, respectively based on the resulting mobility scores from the transgenic strain. To the best of our knowledge, this score exceeds the threshold reported in previous studies, where a z-factor above 0.3 is generally considered feasible for quantitative HTS (21).

### Determining the toxicity of common vehicles for compound screening

Common organic solvents, including ethanol and dimethyl sulfoxide (DMSO), are widely used as co-solvents for compound screening (40). Previous studies have shown that exposure to ethanol and DMSO affects *C. elegans* locomotion, lifespan, and signaling pathways (41–43). To minimize such solvent-related interference in future chemical screening experiments, we evaluated the effects of different dosages of ethanol and DMSO. Based on our observation, worms can tolerate up to 0.4% (v/v) of DMSO (Figure 5a) and up to 1% (v/v) ethanol (Figure S7) without a significant change in their movement. For long-term chemical or pharmaceutical screening, we recommend starting with approximately 70 worms to ensure continuation of the experiment even in the presence of toxicants.

**Figure 5.**
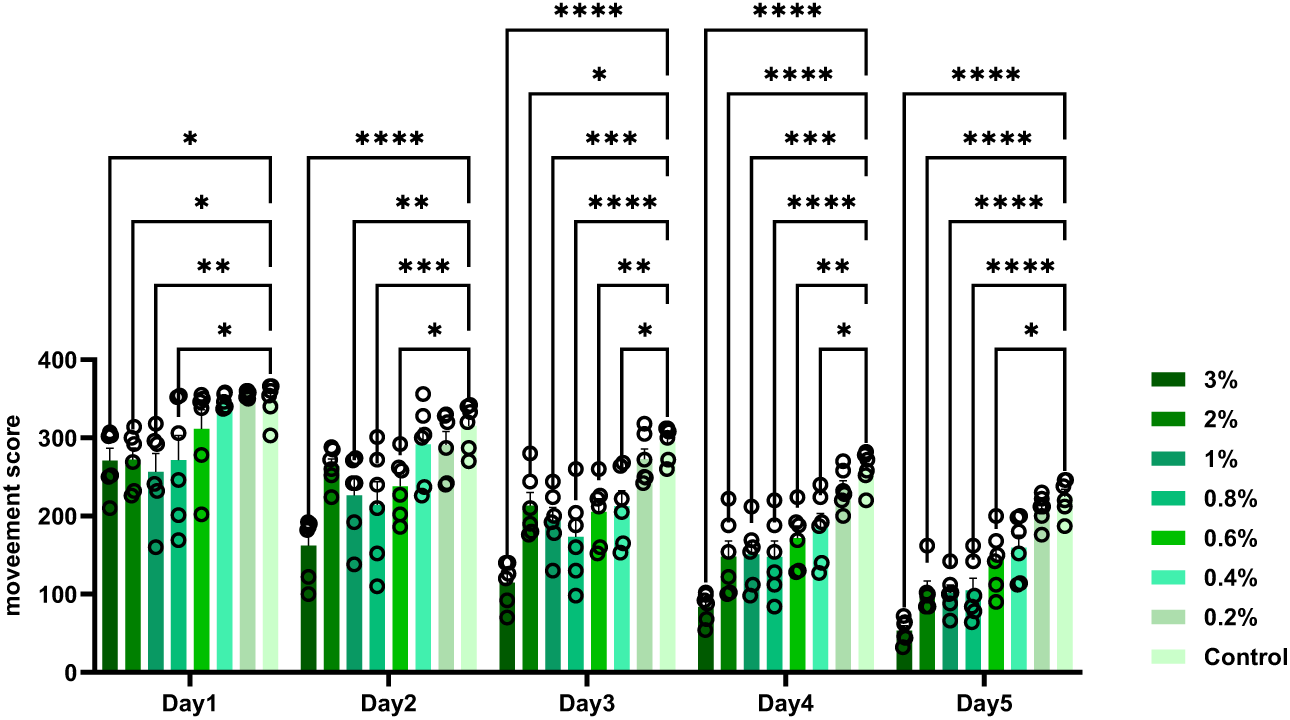
Worms in liquid culture are sensitized to high concentrations of DMSO. Movement comparison between different DMSO dosages in liquid culture,. For all experiments, N = 3, and about ∼70 worms were tested on each trial. For statistical analysis, a two-way analysis of variance (ANOVA) with Tukey’s post-hoc test was used. *P ≤ 0.05, **P ≤ 0.01, ***P ≤ 0.001, ****P < 0.0001.

## Conclusion

This study develops a robust pipeline for HTS with *C. elegans* in liquid culture, by addressing key challenges in worm maintenance, environmental stress, and experimental conditions. This developed framework yielded reliable, reproducible results, with a Z-factor above 0.7. To the best of our knowledge, this is the highest that ever reported for *C. elegans* screening. Our findings also highlight the importance of using compatible utensils when conducting experiments, as well as of experimental parameters such as liquid culture volume, food-to-medium ratio, and life stage selection. Additionally, the comparison between liquid and solid agar culture validated the viability of liquid culture for HTS, with no significant differences observed in worm lifespan or physical characteristics during adult stages up to day 12. In conclusion, our optimized method overcomes the limitations of existing techniques, ensuring minimal chemical and environmental interference while enhancing the reliability of HTS experiments in *C. elegans*. This approach provides a scalable and reproducible platform for a wide range of genetic, metabolic, and pharmaceutical studies, further solidifying *C. elegans* as a versatile model organism in biomedical research.

## Limitations and future directions

While this study optimized conditions to extend the survival of *C. elegans* in liquid culture up to 12 days with no chemical or genetic interference, this is still shorter than their lifespan on solid agar media. This limitation is due to the natural loss of worms during aging and filtering. This may hinder studies focusing on late-life stages or long-term experiments. A possible solution would be to pool the worms from multiple wells to continue the studies. In addition, our pipeline is currently limited to 96-well plates, restricting scalability for higher-throughput applications such as 384-well plates. It should also be noted that because this data acquisition method relies on worm motility, it cannot be applied in cases where chemical and/or genetic interventions do not alter locomotion

## Acknowledgement

This work is supported by the National Institute of General Medical Sciences (NIGMS), the Maximizing Investigators’ Research Award (R35 GM146983). Ms. Yousefsaber received support from the NIGMS-funded T32 Integrative Pharmacological Science Training Program (T32 GM142521, MPI Dr. Anne Dorrance and Gina Leinninger). Dr. Brian Johnson received support from the National Institute of Environmental Health Sciences (NIEHS, R00 ES028744). The authors also thank Drs. Ali Ghiaseddin, Erika Lisabeth, Brian Ackley and Ms. Jennifer Hinman for providing valuable feedback and engaging in discussions that helped refine the ideas presented in this paper. The authors also thank Dr. Patricia K. Dranchak for sharing the ghost bacteria strain that was used in this study with us.

