## supplemental Information for "A Robust Chemical-free Platform for Age-Synchronized *Caenorhabditis elegans* Populations Maintenance for High-Throughput Screening in Aging Studies"

### Supporting Figures and Tables

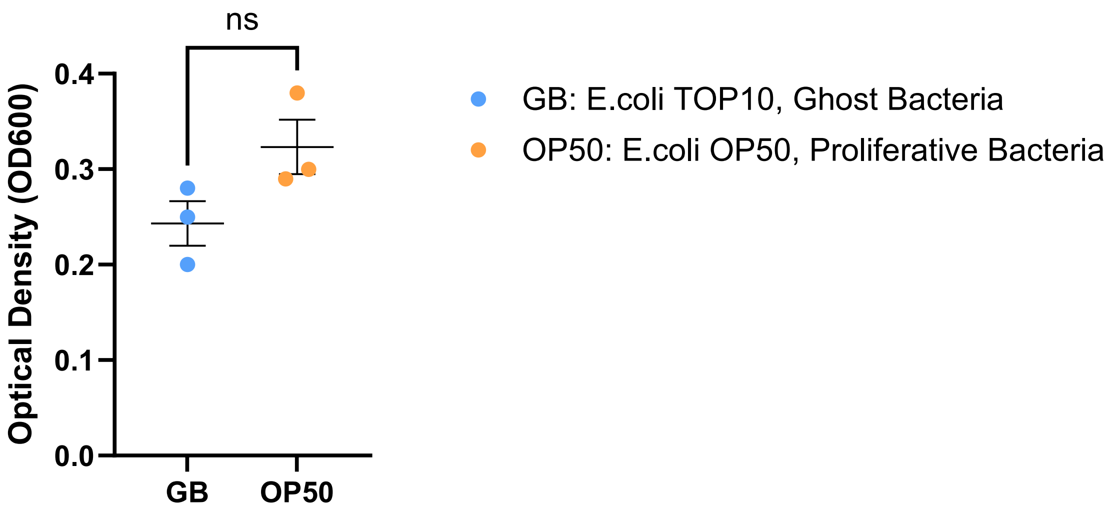


Figure S1. Comparison of optical density (OD600) between *E. coli* TOP10 (Ghost Bacteria) and *E. coli* OP50 (Proliferative Bacteria) after harvesting at 4 °C. Both bacterial preparations reached comparable OD values, confirming reproducibility and suitability across strains as a nematode food source.

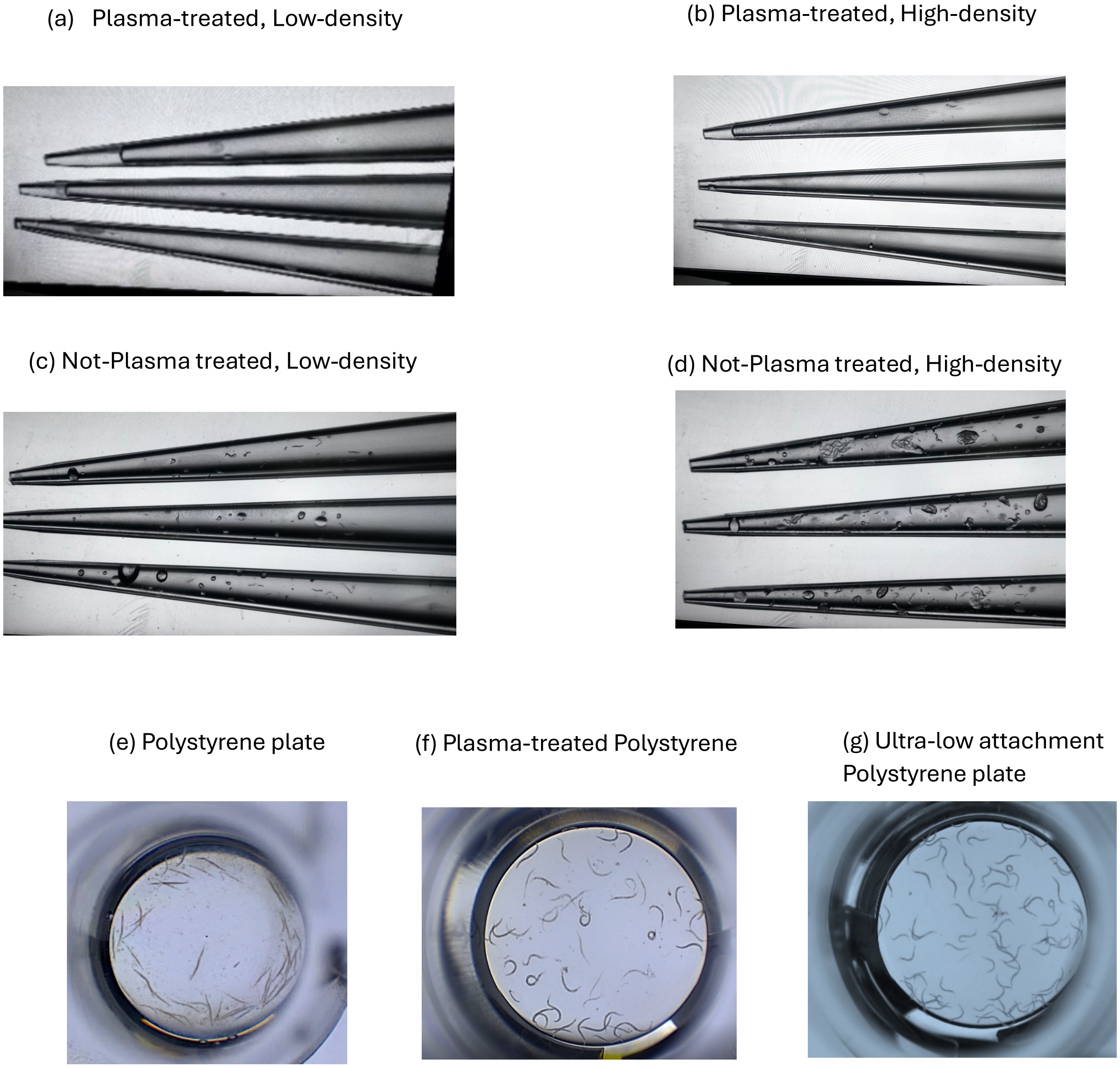


Figure S2. Plasma treatment inhibits worms binding to pipette tips and multi-well plates. (a-d) Oxygen plasma treatment on plastic pipette tips, significantly reduces *C. elegans* attachment on the tip surface. Low-density: ~ 500 worms in 2 mL S-basal solution, High-density: ~ 500 worms in 0.5 mL S-basal solution. (e-g) worms stuck at the bottom of the untreated polystyrene multi-well plates and die (Video S1) while they survive in ultra-low attachment or plasma-treated polystyrene plates (Video S2).

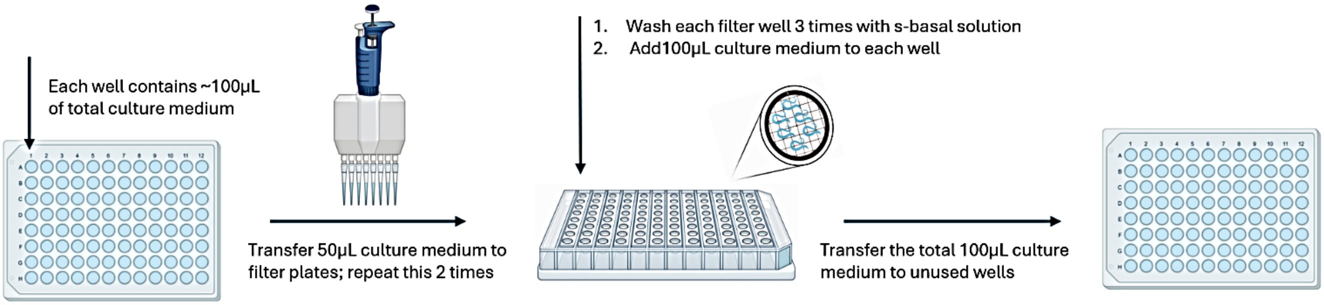


Figure S3. The double step aspiration decreases the chance of transferring bacterial clogs or other contaminants to unused wells.

(a) (b)

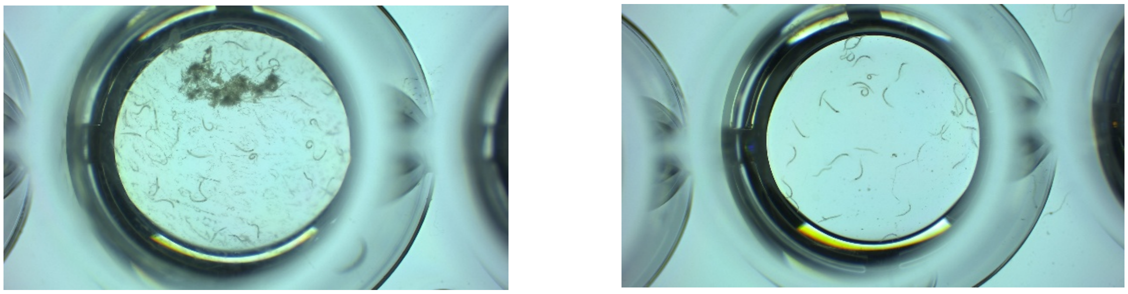

Figure S4. Mesh filters aid with removing bacterial clogs from each well. (a) Before filtering, (b) after filtering. Daily usage of mesh filters with 20 µm pore size not only separate adult population from their offsprings but also remove the bacterial clogs.

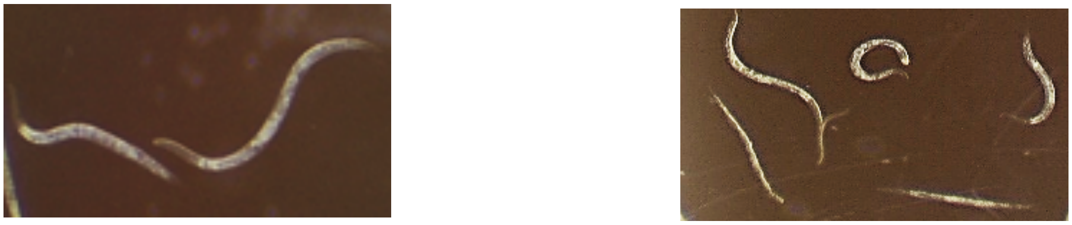

Worms in the presence of cholesterol

Worms without cholesterol in their growth media

Figure S5. Worms' morphology and size is highly impacted by the presence of cholesterol in liquid media. Removing cholesterol leads to starvation; where they get thinner and do not more than 6 days in liquid medium.

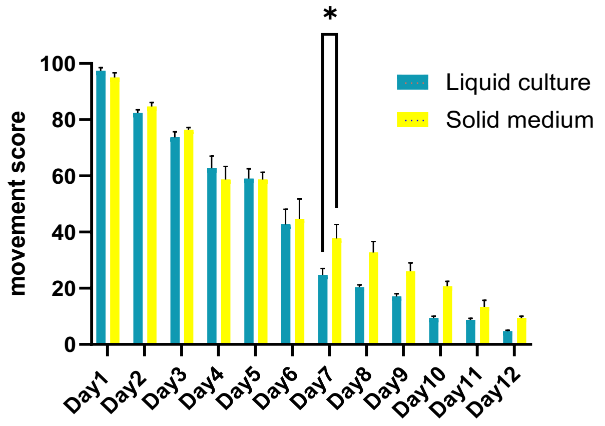


Figure S6. The movement score obtained with Wmicrotracker from liquid and solid culture are consistent. Analysis of *C. elegans* movement in liquid and solid culture to validate results. For statistical analysis, a two-way analysis of variance (ANOVA) with Tukey's post-hoc test was used. *P ≤ 0.05, **P ≤ 0.01, ***P ≤ 0.001, ****P < 0.0001.

(a)

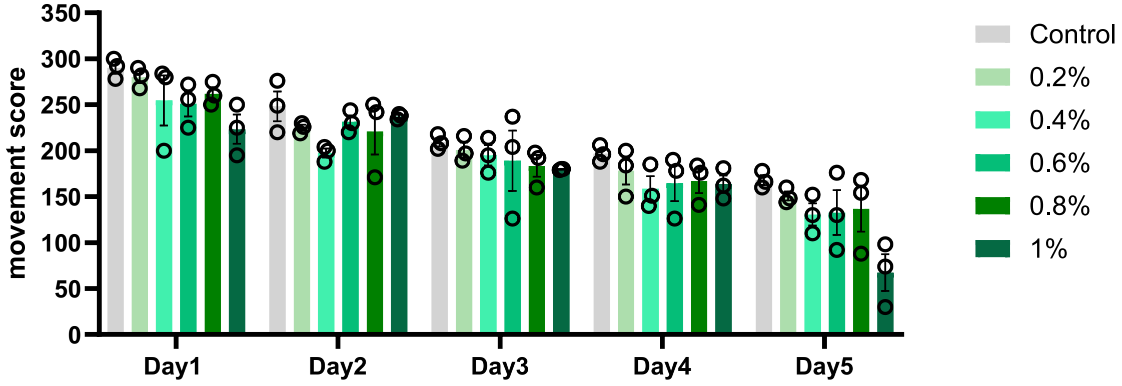


(b)

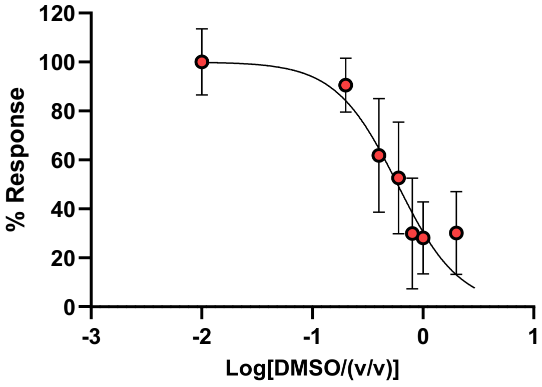

Figure S7. Worms in liquid culture tolerate up to 1% ethanol (v/v) in liquid culture. Movement comparison between different ethanol dosages in liquid culture, (b) Dose response curve: the effect of different DMSO concentrations on movement deficits on day 5 adulthood. For all experiments, N = 3, and about ~70 worms were tested on each trial. For statistical analysis, a two-way analysis of variance (ANOVA) with Tukey's post-hoc test was used. *$P \leq 0.05$, **$P \leq 0.01$, ***$P \leq 0.001$, ****$P < 0.0001$.

#### Image analysis:

Our optimized HTS method is based on movement scores obtained using the Wmicrotracker system, which measures infrared beam interruptions as an indicator of locomotor activity. However, it is important to note that variations in the number of worms per well directly affect mobility scores. Therefore, incorporating worm counts is essential, particularly when comparing different transgenic strains or compound treatments over multiple days, as is often required in aging studies. To address this, we propose an image-based approach using ImageJ for worm quantification and subsequent data normalization.

Here we provide the customized ImageJ macro script used in this study (Figure S8.a). Brightfield images of 96-well plates were acquired with a microscope camera at 2X magnification, ensuring full coverage of each well (~840 pixels in diameter, corresponding to ~7.9 mm). All images were labeled consistently to align with the identifiers used in the movement score datasets, thereby facilitating integrated statistical analysis. Prior to running the macro, directory folders must be created manually, including an input folder for images and an output folder containing two subfolders ("overlay" and "summary"), as ImageJ does not generate folders automatically. Before batch processing, it is recommended to first analyze a single image to confirm appropriate parameter settings, with particle size set to 40–5000 pixels$^2$ and circularity to 0.0–0.4.

As shown in Figure S8, automated worm counts were strongly correlated with manual counts ($R^2 = 0.89$, $p < 0.0001$). Linear regression yielded a slope of 0.97 (95% CI: 0.91–1.02), indicating no significant proportional bias and only a modest offset at low counts. Bland–Altman analysis demonstrated a mean bias of –0.68 worms, with 95% limits of agreement ranging from –10.16 to +8.80 worms, suggesting no systematic deviation between automated and manual methods. These findings confirm that the automated pipeline provides accurate estimates across a wide range of worm densities. Accordingly, automated worm counts can be reliably incorporated for normalization of mobility scores in high-throughput screening applications. Moreover, compared to thrashing-derived N2/Tau ratios (Day 3 = 1.9, Day 5 = 2.2), raw Wmicrotracker mobility scores overestimated strain differences (Day 3 = 2.8, Day 5 = 3.0). Normalization by manual worm counts improved agreement (Day 3 = 1.4, Day 5 = 1.8), while normalization by automated worm counts yielded closer agreement on Day 3 (1.8 vs 1.9) but underestimated the ratio on Day 5 (1.5 vs 2.2). These results indicate that worm-count normalization reduces bias inherent in raw mobility scores, with automated counts providing a practical alternative to manual counting for large-scale HTS, though some discrepancies remain at later time points.

```
// Set input and output folders
inputDir = "C:/Users/MSU/HTS Worms/Input_images/";
outputOverlay = "C:/Users/MSU/HTS Worms/Results/Overlay/";
outputCSV = "C:/Users/MSU/HTS Worms/Results/Summary/All_Results.csv";

// Get list of all .jpg files
list = getFileList(inputDir);
setBatchMode(true);
run("Clear Results");

for (i = 0; i < list.length; i++) {
    if (endsWith(list[i], ".jpg")) {
        open(inputDir + list[i]);
        title = getTitle();
        name = replace(title, ".jpg", "");

        run("8-bit");
        run("Auto Threshold", "method=Triangle ignore_white");
        run("Watershed");
        run("Fill Holes");

        // Append results, do not clear
        run("Analyze Particles...", "size=40-Infinity circularity=0.00-0.35 show=Overlay display exclude include summarize add");

        saveAs("PNG", outputOverlay + name + ".png");

        close();
    }
}

// Save all accumulated results to a single CSV file
saveAs("Results", outputCSV);
```

(a)

(b)

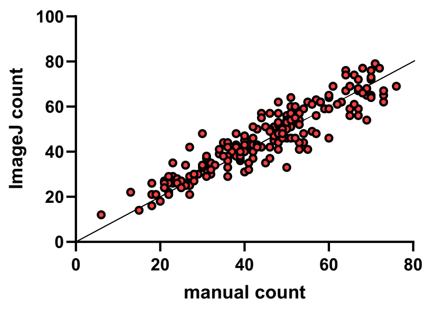


(c)

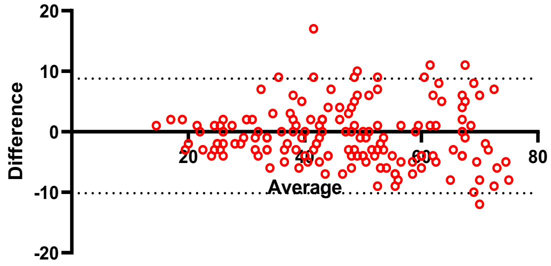


(d)

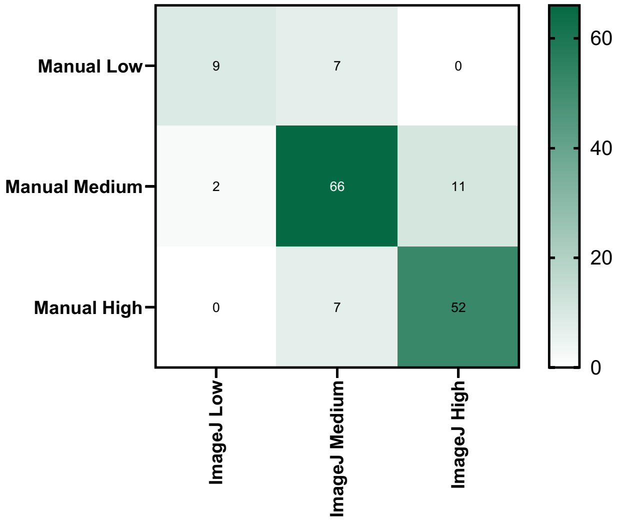


| |
|---|
| Number of observed agreements: 127 (82.47% of the observations) |
| Number of agreements expected by chance: 66.3 (43.06% of the observations) |
| Kappa= 0.692<br>SE of kappa = 0.053 |
| Kappa between 0.61 and 0.80: Substantial agreement |

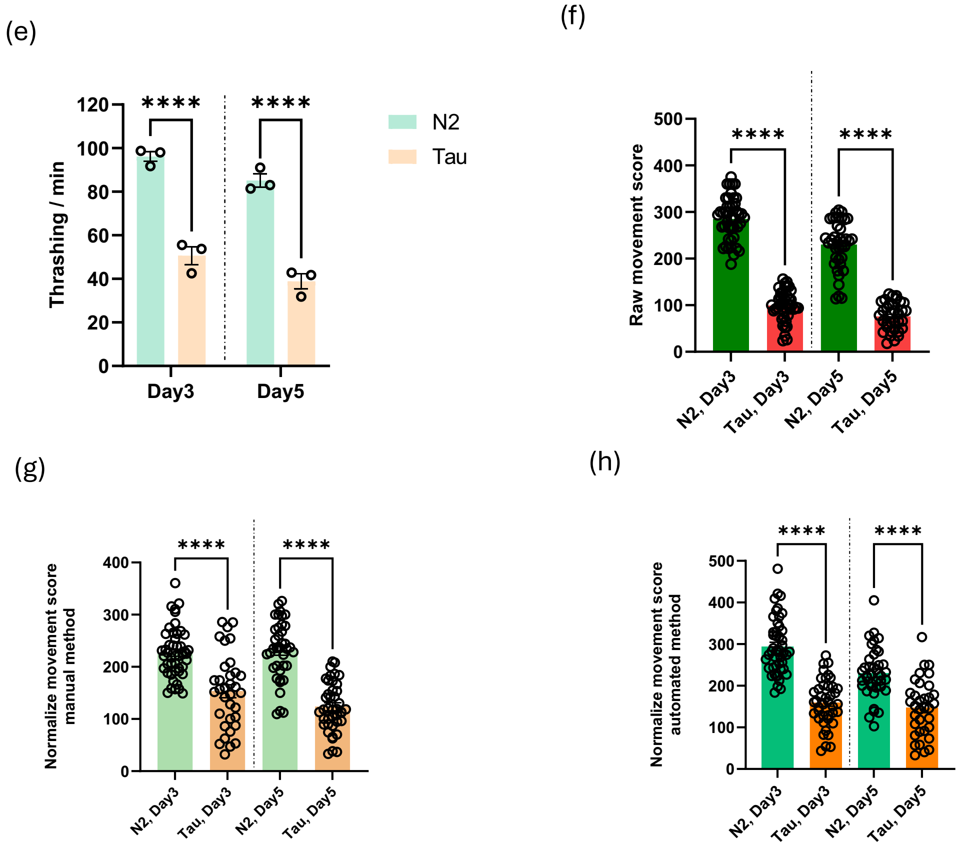


Figure S8. Image analysis makes the statistical analysis faster, reliable and reproducible. (a) Macro script to use in Image J software. The directories should be adjusted based on the specific folder and file location. (b) The correlation between manual and automated counts. Linear regression suggests no significant proportional bias and only a modest offset at low counts. (c) Bland–Altman analysis. The analysis indicates no systematic deviation between automated and manual methods. (d) Heatmap of agreement between manual worm counts and ImageJ-based counts across three categories (Low = 0–25, Medium = 26–50, High = 51–75). Each cell shows the number of images classified into a given category by manual scoring (rows) versus ImageJ (columns). Darker diagonal cells indicate strong agreement between methods, while off-diagonal cells reflect instances of under- or over-estimation by ImageJ relative to manual counts. (e-h) Comparison of N2/Tau locomotion across methods. While raw Wmicrotracker mobility scores overestimated the ratio on day 3 and 5, normalization by manual counts or automated counts improves agreement.
